# Forecasting dominance of SARS-CoV-2 lineages by anomaly detection using deep AutoEncoders

**DOI:** 10.1101/2023.10.24.563721

**Authors:** Simone Rancati, Giovanna Nicora, Mattia Prosperi, Riccardo Bellazzi, Marco Salemi, Simone Marini

## Abstract

The coronavirus disease of 2019 (COVID-19) pandemic is characterized by sequential emergence of severe acute respiratory syndrome coronavirus 2 (SARS-CoV-2) variants, lineages, and sublineages, outcompeting previously circulating ones because of, among other factors, increased transmissibility and immune escape. We propose DeepAutoCoV, an unsupervised deep learning anomaly detection system to predict future dominant lineages (FDLs). We define FDLs as viral (sub)lineages that will constitute more than 10% of all the viral sequences added to the GISAID database on a given week. DeepAutoCoV is trained and validated by assembling global and country-specific data sets from over 16 million Spike protein sequences sampled over a period of about 4 years. DeepAutoCoV successfully flags FDLs at very low frequencies (0.01% – 3%), with median lead times of 4-17 weeks, and predicts FDLs ∼5 and ∼25 times better than a baseline approach For example, the B.1.617.2 vaccine reference strain was flagged as FDL when its frequency was only 0.01%, more than a year before it was considered for an updated COVID-19 vaccine. Furthermore, DeepAutoCoV outputs interpretable results by pinpointing specific mutations potentially linked to increased fitness, and may provide significant insights for the optimization of public health *pre-emptive* intervention strategies.

**Key Points:**

1. **Introduction of DeepAutoCoV:** The article introduces DeepAutoCoV, an unsupervised deep learning anomaly detection system designed to predict future dominant lineages (FDLs) of SARS-CoV-2. FDLs are defined as viral (sub)lineages that will constitute more than 10% of all viral sequences added to the GISAID database in a given week;
2. **Performance and Predictive Capability**: DeepAutoCoV successfully flags FDLs at very low frequencies (0.01% to 3%), with median lead times of 4 to 17 weeks before they become dominant. It predicts FDLs approximately 5 to 25 times better than baseline approaches. For instance, the B.1.617.2 vaccine reference strain was identified when its frequency was only 0.01%, over a year before it was considered for vaccine updates;
3. **Interpretable Results and Mutation Identification**: The system provides interpretable results by pinpointing specific mutations that may be linked to increased fitness, offering insights that can optimize public health interventions. Key FDL mutations, such as those found in Delta and Omicron variants, are identified and analyzed for their potential impact on viral spread and immune escape;
4. **Advantages and Applications**: DeepAutoCoV is advantageous because it does not require prior assumptions about which protein sites are more likely to mutate. Its application in genomic surveillance systems could significantly reduce the time needed for public health responses to emerging variants, enabling early interventions such as vaccine updates;
5. **Evaluation and Comparisons**: The performance of DeepAutoCoV was tested over four years of global and national surveillance data, demonstrating superior predictive power compared to other supervised and unsupervised methods. The system is periodically updated to adapt to the evolving viral landscape, making it a robust tool for ongoing surveillance efforts.

## Introduction

The COVID-19 pandemic has exerted an unprecedented impact on global health, economy, and daily life causing nearly 7 million deaths between December 2019 and January 2024 [1]. Since its first introduction in our species, the original SARS-CoV-2 Wuhan strain [2], [3] has rapidly diversified in distinct (sub)lineages classified by specific genotypic mutations in the Spike glycoprotein [4], [5], [6]. The Centers for Disease Control and Prevention (CDC) has also designated several variants being monitored (VBM) and variants of concern (VOC) – e.g., Alpha (B.1.1.7), Beta (B.1.351), Gamma (P.1), Delta (B.1.617.2), or the currently circulating Omicron (B.1.1.529) and their (sub)lineages – associated with increased transmissibility, more severe disease (e.g., increased hospitalizations or deaths), significant reduction in neutralization by antibodies generated during previous infection or vaccination, or reduced effectiveness of treatments or vaccines [7], [8]. Novel SARS-CoV-2 (sub)lineages capable of outcompeting the ones in circulation have been attributed to the occurrence of secondary, tertiary, quaternary, and quinary COVID-19 epidemic waves [9]. Thanks to genomic surveillance systems, like the one successfully implemented in Africa [10], hypermutated strains under heavy immune pressure, e.g., the emerged JN.1 [10], are nowadays recognized and labelled as VBM or VOC. The World Health Organization (WHO) has outlined specific criteria to select vaccine reference strains in the context of circulating viruses [11]. Unfortunately, as hundreds, if not thousands SARS-CoV-2 (sub)lineages continue to emerge, it is unfeasible to conduct experimental risk assessments or predict which ones will gain dominance in any area of the world; a practical limit as to how often vaccine composition changes need to be implemented, either regionally or globally [12].

Artificial Intelligence (AI), especially deep learning, has been used for several purposes: i) to predict SARS-CoV-2 mutations at specific amino acid sites of the Spike protein [13], [14], [15], [16]; ii) to characterize drivers of emerging strains evolution by calculating antibody escape and binding affinity of the protein to ACE2, which allows viral entry in host cells [17] and iii) to classify sequenced viromes into known (sub)lineages. However, a fundamental limit of current approaches is the inability to identify *novel* emerging (sub)lineages that will spread in the future. This is a critical task in genomic surveillance as it would allow a rapid deployment of countermeasures. To help bridging this gap we propose DeepAutoCoV, a novel approach to forecast future dominant SARS-CoV-2 (sub)lineages (FDLs) based on anomaly detection via unsupervised learning [18], [19]. We define an FDL as any (sub)lineage that will constitute a sizable percentage (10% or more) of all the Spike proteins sequenced over a specific time interval (one week). DeepAutoCoV main goal is to predict (discover) FDLs as soon as possible, i.e., when they are still low-frequency, in order to design proper countermeasures before they spread and become dominant. Our assumption is that, given a steady pathogenic landscape represented by all circulating (sub)lineages, an FDL is detectable as an anomaly carrying specific mutations, which makes them fitter and, therefore, likely dominant in the future. Being an unsupervised anomaly detection system, the key advantage of DeepAutoCoV is that it does not require any *a priori* assumption on which protein sites or domains are more likely to mutate to generate FDLs.

We tested DeepAutoCoV over ∼4 years of global and national surveillance using real-world data. DeepAutoCoV training strategy ensures that the system is updated periodically, which is critical in an evolving viral landscape. Our simulation indicates that using DeepAutoCoV in a genomic surveillance system could help substantially reduce the time required for deployment of public health countermeasures based on epidemiology and experimental data.

## METHODS

### Definition of Future Dominant Lineages (FDLs)

We define FDL as any (sub)lineage reaching or exceeding a 10% threshold frequency among all viromes sequenced in a single week. The threshold is user-defined and can be changed to reflect epidemiology and/or public health considerations. FDLs are calculated independently for each data set (see below).

### DeepAutoCoV principles

So far, work on SARS-CoV-2 (sub)lineages and mutation prediction has been focused on using AI to label new sequences according to a list of known (sub)lineages, as a supervised multi-class problem [6], to evaluate the probability of becoming VOC [16], [20], to predict mutations likely to arise [14], [15], [16], [17] or to rank them for immune escape and fitness potential [16]. Herein, our task is *fundamentally different*, and based on unsupervised anomaly detection, i.e., given a new sequence, we aim to predict whether it is an FDL or not, i.e. if it belongs to a future (sub)lineage that will eventually become dominant.

### Data sets

We downloaded all Spike protein sequences from GISAID, along with their Pango Lineages [6], sampled between December 24^th^, 2019, and November 8^th^, 2023 (total 16,187,950 proteins in 198 weeks).

We focused on the Spike protein as it is the target of antibody-mediated immunity and the primary antigen in current vaccines [21]. Furthermore, by considering only the Spike protein at the amino acid level instead of, for example, the whole genome at the nucleotide level, we greatly reduced the feature space (see “Feature encoding, ground truth, and outcomes”, below). To assure high quality we retained only Spike proteins without missing or unrecognized amino acids; and with lengths *l* ± 30 amino acids (aa), where *l* is the median length of the corresponding lineage (bounds between 1241 and 1301 aa).

We kept the last attributed label in temporal order for identical sequences with inconsistent labels. Upon manual inspection, we removed all Omicron proteins with a collection date that predates November 2021. After filtering (total 4,506,983 proteins), we generated five data sets: four representing nations of interest (USA, UK, France, and Denmark) to simulate national genomic surveillance; and one that includes all sequences regardless of the country of origin, representing a global surveillance program (**Table S1**). The four nations were selected to assess the robustness of our AI model’s ability of FDL early identification in settings with both high (USA and UK, 1,294,936 and 962,496 sequences after filtering) and low (France and Denmark, 140,757 and 312,612 sequences after filtering) sequence volumes.

### Feature encoding, ground truth, and outcomes

We converted aa sequences into fixed-length numeric features using *k*-mers, with *k* = 3 according to previous works [22], [23]. In other words, we represent each Spike protein as a binary vector of fixed length where each slot corresponds to the presence (1) or absence (0) of a 3-aa *k*-mer. If DeepAutoCoV flags an FDL before it reaches the 10% threshold, we consider it a true positive. Conversely, we consider as a false positive a non-FDL protein flagged as FDL, or an FDL flagged after the 10% threshold is reached (i.e., when the FDL is no longer “future”, but a *current* dominant lineage).

### Anomaly Detection System

DeepAutoCoV is based on a deep learning AutoEncoder. The AutoEncoder is designed to handle *k*-mers in the input layer. It consists of an encoding layer followed by a series of dense layers, and a dropout layer to reduce overfitting (**Figure 1A**). The decoder mirrors the encoder’s structure, and the output layer reconstructs the input data. We provide a detailed description of the AutoEncoder in Supplementary Methods. Code is available at **https://github.com/simoRancati/DeepAutoCoV**. The AutoEncoder is trained over fifty epochs with a batch size of 256, utilizing the mean square error (MSE) loss function and Adam optimizer, with TensorFlow and Keras Python libraries [24], [25].

**Figure 1.**
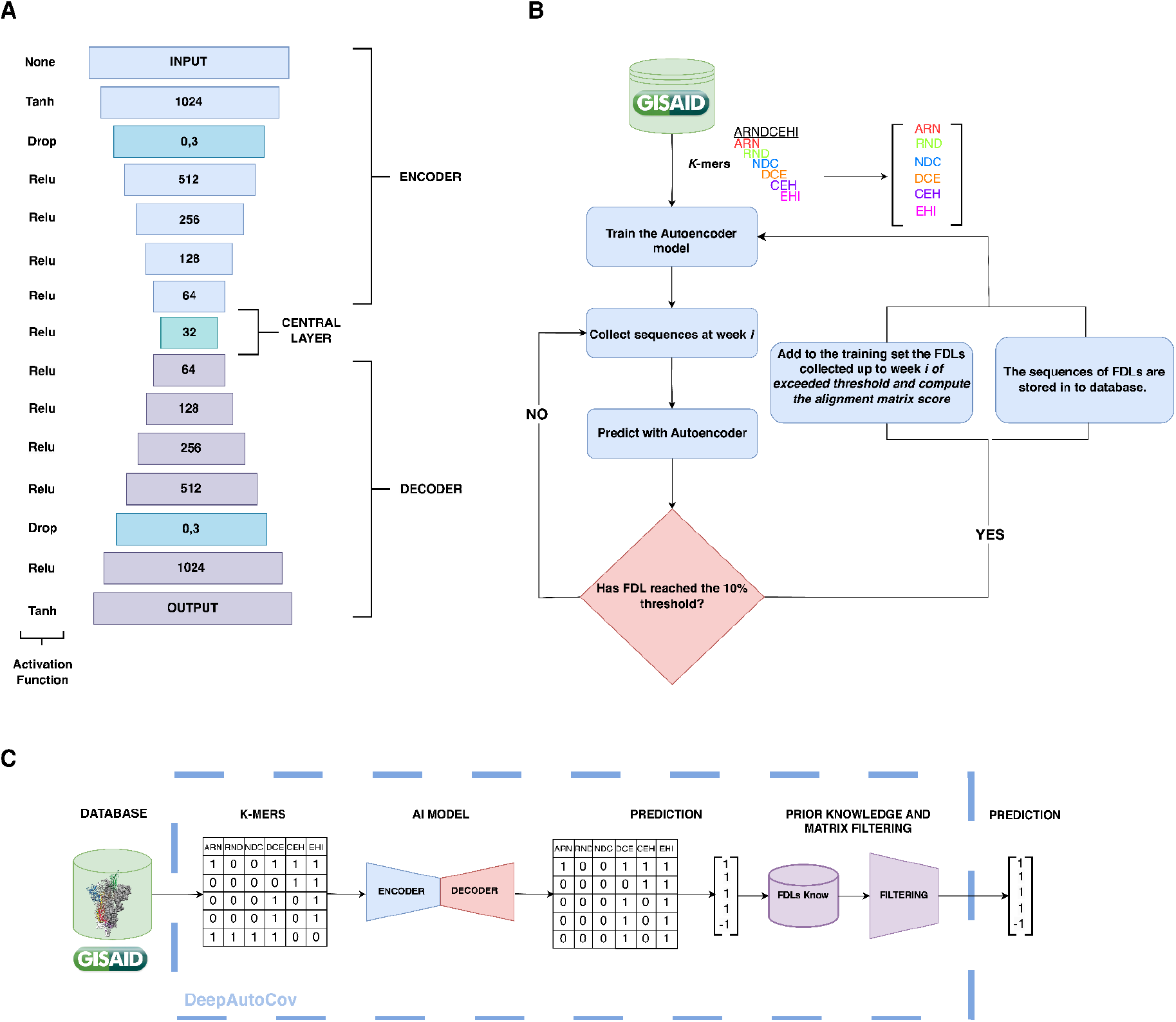
DeepAutoCoV architecture. **A)** Layered architecture with decreasing neuron counts in the encoding layers (from 1024 to 64) and increasing counts in the decoding layers (from 64 to 1024). A central layer with 32 neurons introduces Gaussian noise to enhance generalization. The decoding layers mirror the encoding structure. The output layer reconstructs the original input data. **B)** Schematic of the genomic surveillance simulation. SARS-CoV-2 Spike protein sequences are gathered, converted into *k*-mers, and filtered. An AutoEncoder outlier detection system is trained. Each week, newly uploaded sequences are evaluated as either FDLs or not. When a FDL reaches the 10% threshold, the model is retrained. **C)** Overview of DeepAutoCoV process: *k*-mers extraction, AutoEncoder, prediction and filtering.

Anomalies are detected by identifying data points that deviate significantly from their learned reconstructions (conventionally over 1.5 standard deviations from the median [26], [27]), based on the Mean Squared Error (MSE).

After a target protein is flagged as FDL, we perform additional filtering by scoring via Point Accepted Matrix (PAM) [28]. First, we compute the average PAM score between all the training set FDL proteins and the original Wuhan sequence. If the PAM score between the target protein and the original Wuhan sequence exceeds the average score, we assume the target sequence is not an anomaly. We also automatically ignore proteins that are exactly identical to proteins included in the training set, and as such, non-FDL by definition.

### Simulation of genomic surveillance and FDL emergence

We emulate the application of DeepAutoCoV within genomic surveillance at a global and national level (**Figure S1**). We assume that sequences are uploaded and analyzed weekly by DeepAutoCoV.

We initially trained DeepAutoCoV with the sequences collected during the first week (**Table S8**). Note that the initial training set is different for each dataset, and represents the viral steady-state of the corresponding area. To reduce the number of features (*k*-mers), we keep only *k*-mers that are present in at least 1% of the sequences in the first training set. Once the AutoEncoder is trained, Spike proteins collected in the following period, with a 1-week pace, are processed for anomaly detection. When an FDL reaches the dominance threshold (10%) in a given week, the FDL anomaly detection process halts, and the model is re-trained and the average PAM score is recalculated by including all the proteins belonging to the newly emerged FDL proteins that have appeared so far (**Figure 1B, 1C**).

Performances are evaluated weekly over the top 100 proteins based on MSE ranking. The choice of considering 100 sequences instead of the whole weekly batch is motivated by emulating a realistic scenario, where DeepAutoCoV is the first line in an integrated surveillance program (**Figure S1**).

Realistically, over hundreds or thousands of collected proteins per week, only a fraction can be thoroughly investigated *in vitro*. For each data set, the model is evaluated in terms of (a) lead time, i.e., the number of weeks between first detection of an FDL and the week when it reaches 10% threshold; (b) frequency at week of detection, i.e. the ratio between a specific FDL target proteins and the total proteins collected at the week of first detection; (c) Prioritization, i.e. the ability to rank FDL proteins as top scoring (based on MSE); (d) Positive Predictive Value (PPV), i.e., the fraction of correctly flagged FDLs over the total flagged proteins.

The code to reproduce the simulation is available at **https://github.com/simoRancati/DeepAutoCoV**.

### Model comparisons

We compared the performance of DeepAutoCoV against both supervised and unsupervised methods. Specifically, we developed supervised approaches using Logistic Regression (LR), Spike2Signal (S2S) [29], Spike2vec (S2V) [30]; and an unsupervised linear distance-based (DB) anomaly-detection method, i.e., an anomaly detection model using linear distance instead of the features extracted by the AutoEncoder. As for DeepAutoCoV, the performance of the competing methods are computed over their top 100 flagged (i.e., predicted as anomalies by the specific method) proteins per week. PPV p-values were calculated by sampling n = 100 random Spike proteins per time point, assuming to be FDLs and comparing them to the top 100 Spike proteins selected by DeepAutoCoV (Wilcoxon paired test). The Supplementary Methods provide a detailed description of competing models.

## RESULTS

### DeepAutoCoV flags FDLs with a median lead time of 17 weeks

To calculate the lead time, for each FDL we calculated the difference in weeks between the first true positive detection by DeepAutoCoV of that FDL, and the FDL reaching the 10% threshold (for visual representation of the results, see Figure S2 and Figure S3). In the Global data set, FDLs are flagged with a median lead time of 17 weeks before reaching the 10% threshold, and an interquartile range (IQR) of 11 to 32 weeks. In the USA, UK, Denmark, and France data sets, the respective median lead times are 12 weeks (IQR 7, 37); 6 weeks (IQR 2, 11); 5 weeks (IQR 2, 11); and 11 weeks (IQR 6, 20). The interval between the 1^st^ and the 3^rd^ quartile varies from 30 (USA) to 9 weeks (Denmark and UK) (**Figure 2**). DeepAutoCoV flags FDLs when their frequency is still very low (**Table 1**), ranging from 0.01% (Global data set) to 3% (UK). Per-(sub)lineage details are provided in **Table S7** and **Figure S2, S3**.

**Table 1.** DeepAutoCov results over all data sets.

| Data set | Lead time<br>(weeks)<sup>1</sup> | FDL<br>frequency at<br>discovery<sup>2</sup> | PPV<sup>3</sup> | Prioritization<sup>4</sup> | p-value<sup>5</sup> |
| --- | --- | --- | --- | --- | --- |
| Global | 17<br>(11,32) | 0.1%<br>(0.02%, 0.7%) | 0.30<br>(0.06, 0.42) | 6° place | 4e-06 |
| USA | 12<br>(7,37) | 0.2%<br>(0.05%, 3%) | 0.48<br>(0.10, 0.76) | 3°/4° place | 4e-07 |
| UK | 6<br>(2,11) | 3%<br>(0.1%, 8%) | 0.57<br>(0.24, 0.92) | 3°/4° place | 7e-16 |
| Denmark | 5<br>(2,11) | 0.8%<br>(0.2%, 5%) | 0.49<br>(0.02, 0.90) | 4° place | 4e-03 |
| France | 11<br>(6,20) | 1.5%<br>(0.4% 7%) | 0.47<br>(0.17, 0.70) | 4° place | 7e-12 |

1. Median lead time in weeks (IQR). 2. Median (IQR) FDL frequency at discovery time. Frequency is the proportion of the discovered FDL proteins over the total Spike proteins in the data set in the week. 3. Median (IQR) Positive Predictive Value. 4. Prioritization, i.e., the average ranking position of the first true positive. 5. Wilcoxon paired test (see Methods)

**Figure 2.**
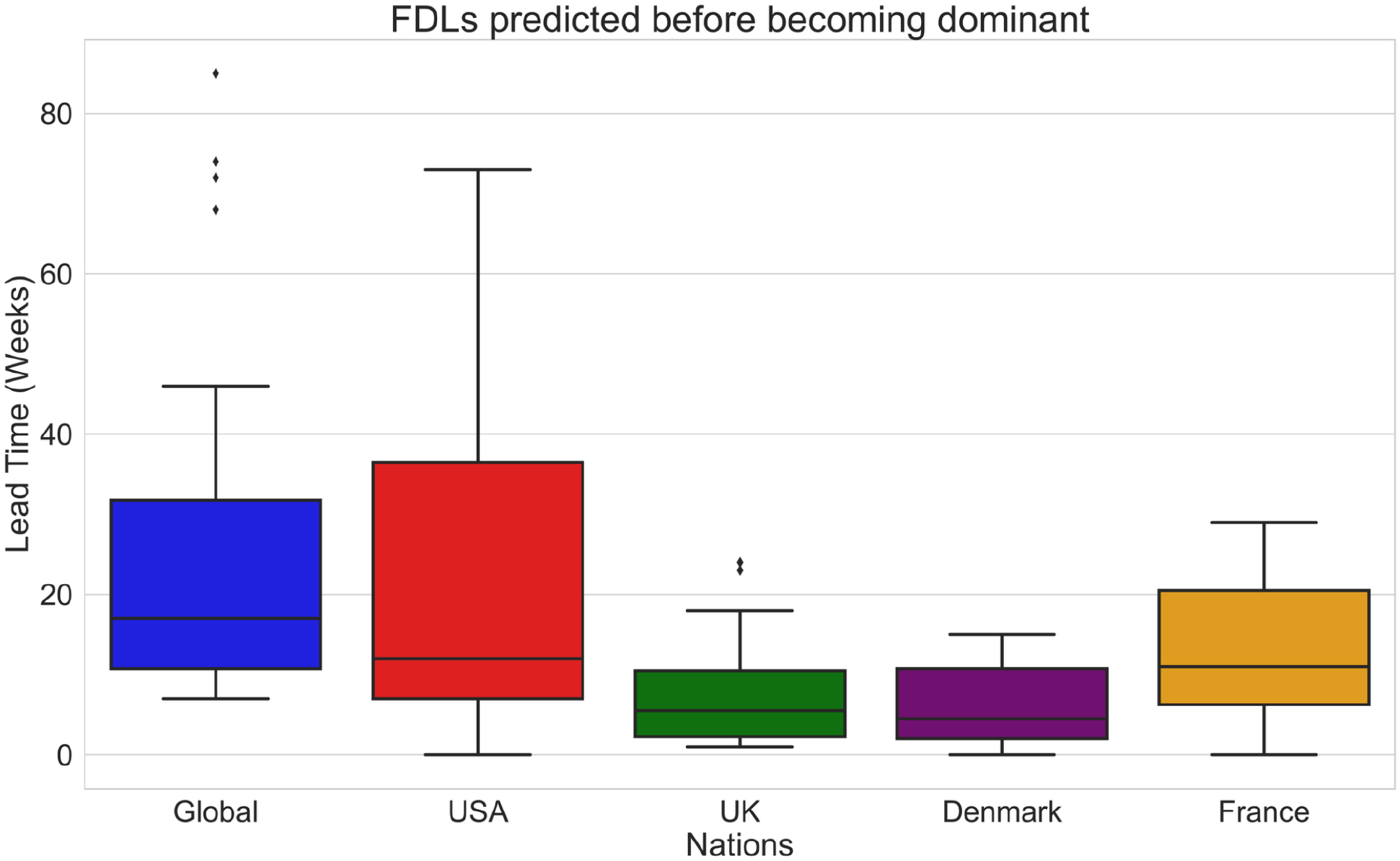
Per-data set DeepAutoCoV performance in terms of lead time (weeks) before reaching the 10% threshold.

### FDL Prioritization

Prioritization is widely used when thousands of instances are evaluated simultaneously [31]. In case of high transmissibility of a virus, it is unfeasible to rapidly test in the wet lab a high number (hundreds to thousands) of sequences collected and flagged as FDL. Therefore, the ability to prioritize the FDL correctly flagged as an anomaly in the top positions is crucial. We evaluate DeepAutoCoV prioritization ability by recording the position of the first FDL correctly identified. For the Global data set, the first true positive is detected on average in the 6th position across weeks. For the nation-specific dataset, the average position of the first positive is the 3^rd^ or the 4th. Supervised methods failed to correctly flag any FDL, therefore do not have a prioritization value.

### FDL Positive Predictive Value

The PPV is the ratio between true positives and the sum of true positives and false positives. DeepAutoCoV median PPV is 0.30 (IQR 0.06; 0.42, *p* = 4e-06) for the Global Data set, 0.48 (IQR 0.10, 0.76; *p* = 4e-07) for the USA Data set, 0.57 (IQR 0.24 0.92; *p* = 7e-16) for the UK Data set, 0.49 (IQR 0.02, 0.90; *p* = 4e-04) for the Denmark Data set, and 0.47 (IQR 0.17, 0.7; p-value 7e-12) for the France data set. Over time (**Figure 3**), PPV moving average follows the same trend in each data set. The total number of proteins for the top ten FDL on each set, along with p-values, is reported in **Figure S4**. PPV (PPV) varies inversely with the size of the data set. The reason can be found by noting the negative correlation (Pearson coefficient: -0.84) between the mean entropy and average PPV across the five datasets – it is easier to detect FDLs when the genomic landscape is more homogeneous. During weeks when most uploaded sequences are unique, flagged FDLs are likely to be false positives. Conversely, when uploaded sequences belong to multiple tight lineage clusters, such as groups of similar strains like the B.1.1.7 (Alpha) lineage or the AY.3 (Delta) sublineage, flagged FDLs are more likely to be true positives.

**Figure 3.**
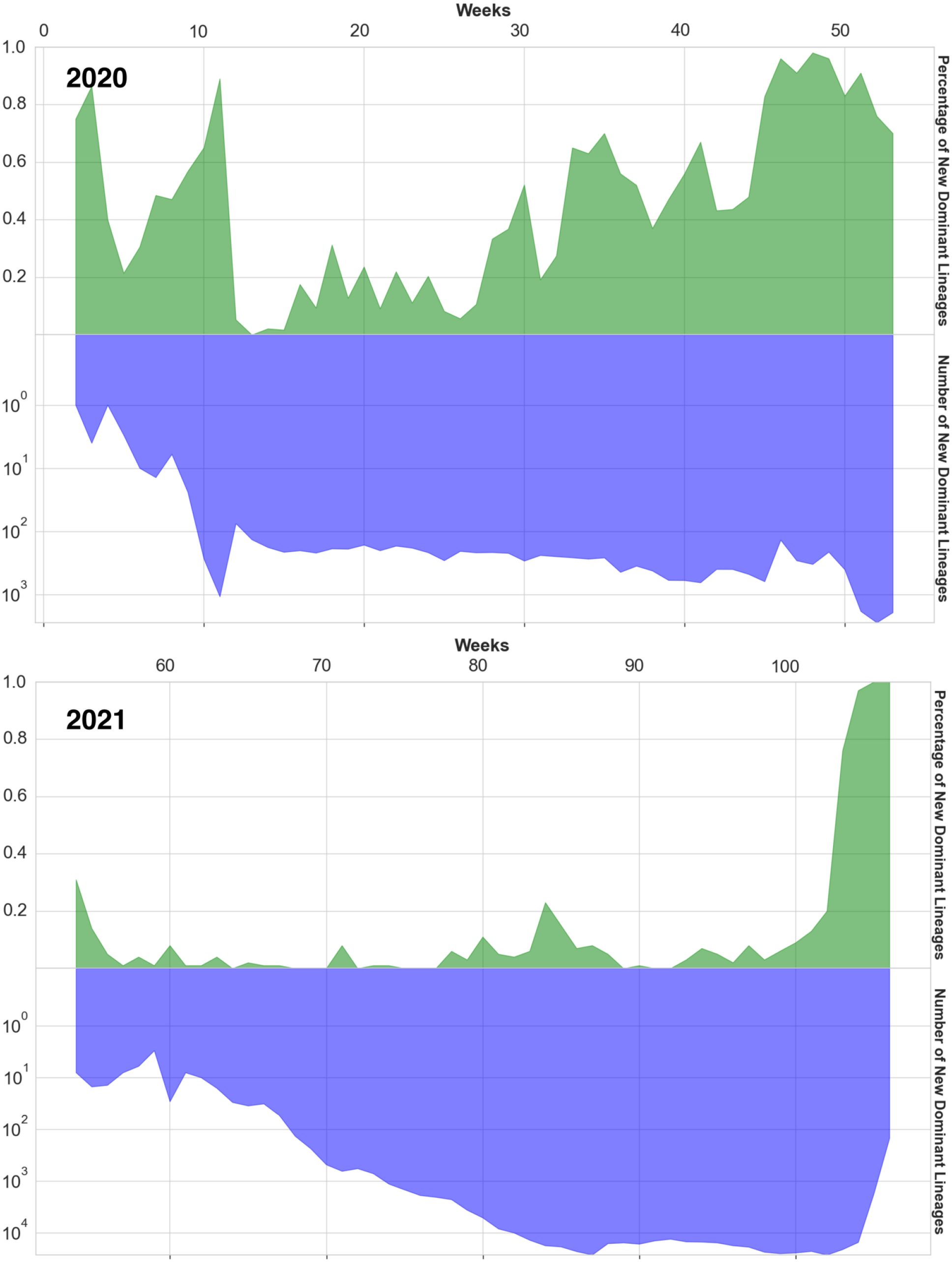
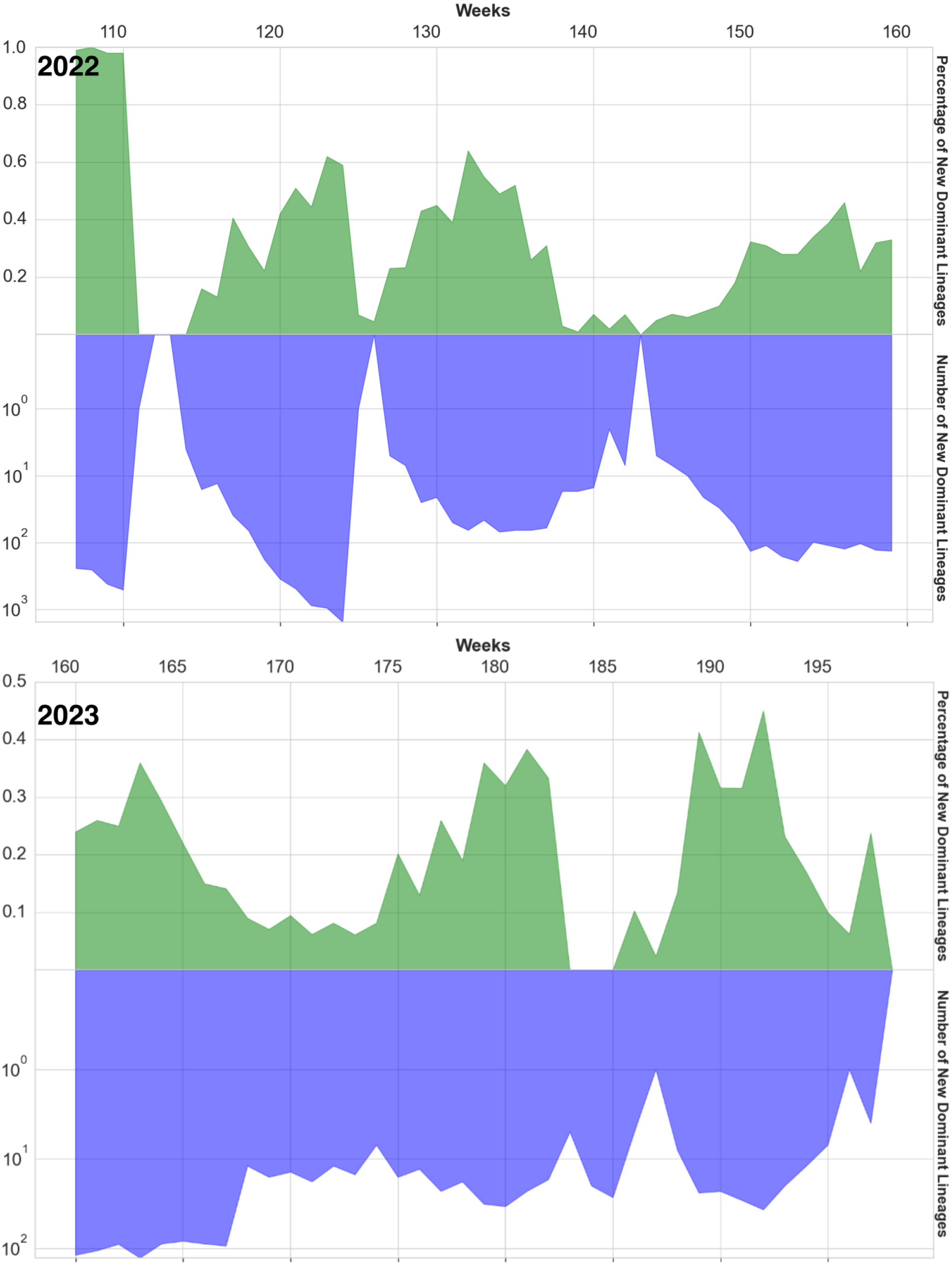
X-axis: simulation weeks; y-axis top side: PPV (green); bottom side: number of new FDL proteins in semi-logarithmic scale, each week (blue).

DeepAutoCoV outperforms the linear approach in the Global data set (median PPV 0.3 and 0.05 respectively). Supervised models do not flag any FDLs (PPV 0), as by design *emerging* FDLs are unknown to the model. Details are provided in **Table S2**.

### DeepAutoCoV predicts the newly emerged KP lineage

We further evaluated DeepAutoCoV on 2,295 newly detected sequences (from 12-01-2023 to 08-18-2024) without retraining the model. Figure S5 shows the top 10 lineages prioritized by DeepAutoCoV. Notably, DeepAutoCoV flagged the newly emerged KP lineage as anomaly and ranked in the very first positions. Currently, the KP lineage is dominating the SARS-CoV-2 landscape: a recent study showed that the KP.2 variant has higher viral fitness and it is characterized by an increased reproduction ability, making it more transmissible than its predecessor, the JN.1 variant [32], [33]. KP.2 has demonstrated a significant capacity to evade immune responses, including those induced by the latest monovalent XBB.1.5 vaccine. This variant includes specific spike mutations (FLiRT mutations) which are believed to enhance its immune escape and transmissibility [32], [33].

## DISCUSSION

### Assumptions and limitations

DeepAutoCoV evaluation is based on the assumption that all the Spike protein sequences are uploaded to the database in the same week they are collected, and that the sampling strategies reflect the actual spread of the virus. We also assume that, when an FDL reaches a frequency of 10% or more among the collected sequences in a given week, we consider it to have emerged, i.e., its diffusion has been noticed through public health data. Focusing on the Spike protein allowed us to significantly reduce the k-mer computational space (k=3). However, this assumption may not hold true when dealing with more complex computational landscapes, such as multiple proteins or whole genomes. These scenarios would likely necessitate greater computational resources or different feature encoding methods, in particular when a possible application to other viruses might be required.

### Advantages and potential applications of DeepAutoCoV

In a genomic surveillance system, DeepAutoCoV would be capable of early flagging hundreds to thousands of FDL proteins, which can be subsequently analyzed in the wet lab, providing a solid guess of which ones will become the next dominant (sub)lineages (**Figures S1, S4**). While this system is designed for SARS-COV-2 Spike proteins, DeepAutoCoV principles such as the simulation framework and the anomaly detection system, could potentially be adapted to other viruses.

Each data set is characterized by different FDLs, i.e. the same (sub)lineage might be dominant in an area but not in another. Furthermore, FDLs common to more than one set might appear at different times, depending on both SARS-CoV-2 epidemiology patterns in specific locales, potential bias (e.g., sampling bias), and filtering of low-quality sequence data (**Tables S3-S7**). For example, Alpha variant (B.1.1.7) strains were detected in UK for the first time in April 2020, but appeared in our filtered data set the 4^th^ week of August 2020, and were flagged as FDL the same week, when their frequency was 0.05%, eleven weeks before the variant reached the 10% threshold (**Table S5**). In the US, the first Alpha variant sequences appeared in the filtered data set during the 1^st^ week of March 2020, and were flagged during the same week, when their frequency was only 0.2%, 52 weeks before they reached the threshold (**Table S4**). DeepAutoCoV performs worse in the Denmark and France data set (**Tables S6** and **S7**). In the Global data set, Alpha was uncovered in the 2nd week of February 2020 and flagged as FDL the same week, when its frequency was 0.7%, 46 weeks ahead of the threshold (**Table S3**). In general, DeepAutoCoV appears to perform consistently well with both high (Global, USA, UK) and low (France, Denmark) sequence volumes. Reflecting the bias in the data sets, neither Beta nor Gamma were found. Despite co-circulating with the Alpha variant, Beta and Gamma infections were limited to regions not well represented in GISAID at that time, such as Brazil [34] and Haiti [35]. The potential usefulness of our approach is also exemplified by its ability to forecast as FDLs several Omicron (sub)lineages (**Tables S3-S7**).

The heavily mutated Omicron variant (B.1.1.529/21K) was first identified by the South African surveillance network at the beginning of the November 2021 4^th^ week^39^, and already designated a VOC by the WHO’s Technical Advisory Group on SARS-CoV-2 Virus Evolution by November 26^th^. Hopefully, molecular monitoring programs will continue to detect and assess hypermutated strains, such as Omicron, at an increasingly fast pace. Yet, several subsequent Omicron sublineages became dominant or co-dominant in different parts of the world. Assessing which ones of all continuously emerging strains might be an FDL, would be unfeasible, if not aided by a prediction algorithm as the one described herein. DeepAutoCoV might also help predicting in advance (sub)lineages that will become vaccine reference strains, such as BA.1. Earliest BA.1 strains were sampled the 1^st^ week of November 2021. In our simulation using the Global data set, BA.1 is detected the same week it appears, when its frequency is only 0.01% and 10 weeks before reaching the threshold. Similarly, BA.5 and XBB.1.5 are flagged 16 (frequency 0.08%) and 23 weeks (frequency 0.1%), respectively, before each variant reaches the threshold. The case of vaccine variant B.617.2 is particularly relevant because it is flagged as FDL April 2020, not only 73 weeks ahead the 10% threshold but also more than one year before Pfizer researchers began to discuss the lineage as a candidate for a vaccine update [36].

The US CDC typically flags a dangerous (sub)lineage after public health data show its concerning capabilities, such as a spike in infections or hospitalizations. Our main purpose is the prediction of FDLs *as early as possible*. In an actual deployment as part of a genomic surveillance system, DeepAutoCoV flagging FDLs ahead of time would allow public health interventions, such as early vaccine design, to potentially prevent the (sub)lineage from spreading to the population. FDLs detection can, therefore, be the first step in a coordinated genomic surveillance effort (**Figure S1**). The next steps after FDL identification would include (a) molecular simulations, i.e., computational techniques modeling the behavior of the Spike protein to study if the FDL structures could provide critical advantages to the virus ability to spread, such as enhanced ability to bind the receptors; and (b) virulence assays or other phenotypic wet lab validation. As resources are limited, it would not be realistic to envision a global or national pandemic surveillance program providing virulence assays or phenotypic wet lab tests to include *all* the sequenced SARS-CoV-2 Spike protein sequences. Importantly, the DeepAutoCoV framework is suitable not only for SARS-CoV-2, but easily generalizable to other rapidly evolving viruses. In a real-case scenario, we could exploit intrinsic characteristics of DeepAutoCoV to fulfill MLOps (ML Operation) principles during deployment. In fact, the MSE patterns could be used to monitor the generalization ability of the model over time, and to evaluate whether model retraining is required. For instance, if MSE becomes systematically higher, we could foresee a potential data drift and a change in the viral population that may require the re-training of the model.

DeepAutoCoV outperforms concurrent approaches, including a 17-week median lead time and top prioritization. The measured PPV ranges from 0.3 to 0.57, depending on the data set. This result should be interpreted in the context of anomaly detection. Differently from a classification task, our goal is not to identify *all* FDL proteins, which given SARS-CoV-2 rapid accumulation of polymorphic sites [37] may be very large and not necessarily useful, but to flag a small number of likely FDL proteins to be further analyzed (Figure S1).

### DeepAutoCoV identifies key FDL mutations

DeepAutoCoV provides interpretable results, i.e., key Spike FDL mutations. Key FDL mutations, in turn, can provide insights about the molecular mechanisms contributing to future dominance. The underlying mechanism revolves around the binary input representation (a sequence of 0s and 1s indicating the absence or presence of specific *k*-mers). Sequences flagged as FDLs can be analyzed by extracting *k*-mers that are wrongly reproduced by the model after compression (**Figure 1A, 1C**). By matching these *k*-mers with the target Spike protein, we can pinpoint critical mutations. For example, while flagging the AY.4 lineage (Delta variant) DeepAutoCoV flags the *k*-mers “NSH”, “SHR”, and “HRR” which occupy positions 680, 681, and 682 respectively. This is indeed a critical position characterizing the Delta variant mutation P681R, which plays a crucial role in enhancing the virus replication and transmission. This mutation is located at the furin cleavage site of the Spike protein, which is essential for the virus ability to enter host cells. The P681R mutation enhances the cleavage of the spike protein into its S1 and S2 subunits, a process that facilitates viral entry into human cells, particularly respiratory epithelial cells. Consequently, this mutation is a key factor that contributed to the Delta variant’s rapid global dominance over [38]. Another example is the BA.2 (Omicron) lineage, where DeepAutoCoV highlights a *k*-mer pinpointing the Y505H mutation. This mutation induces internal conformational changes in the Omicron Spike receptor-binding domain (RBD), leading to a higher binding affinity with the human ACE2 receptor compared to the Delta Spike RBD, ultimately contributing to greater transmission and infectivity. Y505H mutation plays a critical role in stabilizing the RBD-ACE2 complex by mediating an intermolecular hydrogen bond with 98% occupancy, providing a stronger binding affinity [39]. In the 2024-emerged KP lineage, labeled by our system as an anomaly, the FLiRT mutations (R346T, F456L) were pinpointed by DeepAutoCoV as crucial [32], [33].

## Acknowledgments

NIH NIAID R01 AI170187

## Contributions

SM conceived the paper. SM, MS, RB, and MP designed the experiments. SR, GN, and SM, prepared the data and designed the models. SR and GN performed the experiments. All authors analyzed the results and wrote the paper.

