## Supplementary material for "Forecasting dominance of SARS-CoV-2 lineages by anomaly detection using deep AutoEncoders": Supplentary Methods

### Supplementary Methods

#### AutoEncoder model

We converted protein sequences into fixed-length numeric features using *k*-mers as previously described. Briefly, k-mers are substrings of user-defined length k contained in a sequence [22], [23]. For example, given *k*=2, we find in the sequence "ARNDC" the k-mers 'AR', 'RN', 'ND', 'DC'. Each *k*-mer has a boolean value indicating its presence/absence. Smaller *k*-mers are preferred as they limit the computational burden (i.e., smaller number of features, less sparse data). Based on this principle we selected a *k*-mer value of 3. Experiments with k=5 and k=7 provided similar results (data not shown). AutoEncoders are neural networks trained to learn efficient input representations and reconstruct the original data. They consist of an encoder, which compresses the input, a central layer, and a decoder, which reconstructs it. AutoEncoders can detect anomalies by identifying data points deviating significantly from learned reconstructions, for instance using the mean square error (MSE) to calculate reconstruction errors. Data points that have a MSE higher than a 1.5 standard deviations [26-27] are considered anomalous (i.e., FDL proteins).

The AutoEncoder (**Figure 1A**), which is designed to handle *k*-mers in the input layer, consists of an encoding layer with Tanh activation, followed by a series of dense layers with ReLU activation and a dropout layer for reducing overfitting. The central part includes a dense layer with Gaussian noise and 32 neurons. The decoder mirrors the encoder structure. Finally, the output layer reconstructs the input data using the Tanh activation. This architecture choice was motivated by pursuing good performances while ensuring lower training time and costs, as the AutoEncoder is trained on an increasing number of sequences (up to millions). Moreover, when transitioning from a large number of neurons in one layer to a significantly smaller number in the following, we might lose important information necessary for reconstructing the original input accurately, as each neuron in a layer typically represents a feature or a combination of features from the input data; fewer neurons mean fewer features can be captured and represented [40],[41]. For this reason we opted for a smooth decrease (increase) in the number of neurons per layer in the encoder (decoder).

#### Model comparison

To keep training and testing data consistent with DeepAutoCoV, we followed the same strategy described in the Methods section to define training and testing for the competing approaches. Briefly, each week following the baseline data is considered a test set, with FDLs and non-FDLs sequences. When an FDL reaches the 10% threshold, its sequences are added to the training set and the class is switched to non-FDLs, i.e., the (sub)lineage is no longer an anomaly to be detected [38].

*Logistic Regression.* Logistic regression (LR) is one of the fundamental models in machine learning. LR establishes a relationship between a series of independent variables and a binary dependent variable, providing as output the probability that a particular observation belongs to one of the two categories (in our case, Anomaly and non-Anomaly). We used the LR implementation of the Scikit-Learn library (default values).

*Distance-based Model.* This model ranks the sequences based on the distance (k-mer MSE) calculated between the sequences of the *i*-th week, and all the sequences contained in the training set.

*Spike2Vec* is an approach for Spike protein feature extraction based on the *k*-mers frequency and Fourier transform. After embedding proteins into vectors of fixed length, vectors are then mapped into a low-dimensional representation using an approximate kernel method. These vectors are used as an input to machine learning algorithms to identify known lineages via supervised classification. Following *Spike2Vec* original publication [30], we implement a Naive Bayes (NB) approach .

*Spike2Signal* transforms any Spike protein into a signal, i.e. each amino acid is associated to a specific number and a cumulative distribution is calculated. The resulting time series is used to classify known Sars-CoV-2 (sub)lineages using a RESNET deep learning network [29].
