## Supplementary Figures for "Forecasting dominance of SARS-CoV-2 lineages by anomaly detection using deep AutoEncoders"

**Figure S1:** Putative AI-and-genomic-based surveillance approach**. a)** As new sequences are uploaded in genomic databases, the AI model identifies potential FDLs. Sequences flagged as FDLs are further analyzed via molecular simulations and lab testing to confirm their nature (e.g., will become dominant because of enhanced immune escape or transmissibility). **b)** Conventional approach relying on public health data to flag emerging sequences as FDLs and requiring FDLs to spread (e.g., significant surge in cases) before being recognized..

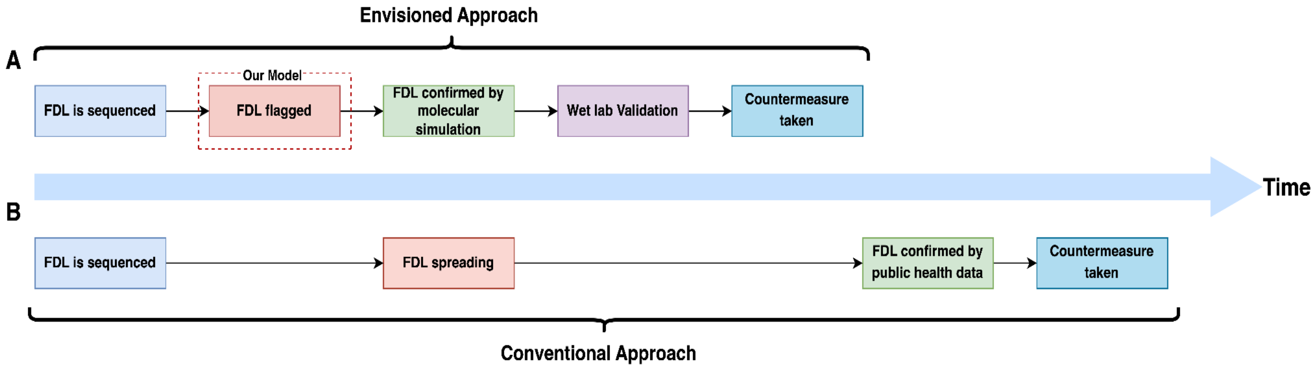

**Figure S2:** DeepAutoCoV **A)** best, and **B)** worst performances for FDL identification on the global data set. The green star indicates the week when the FDL first appears in the data set. The blue square represents the week for first recognition. The red dot indicates the week when the FDL becomes dominant.

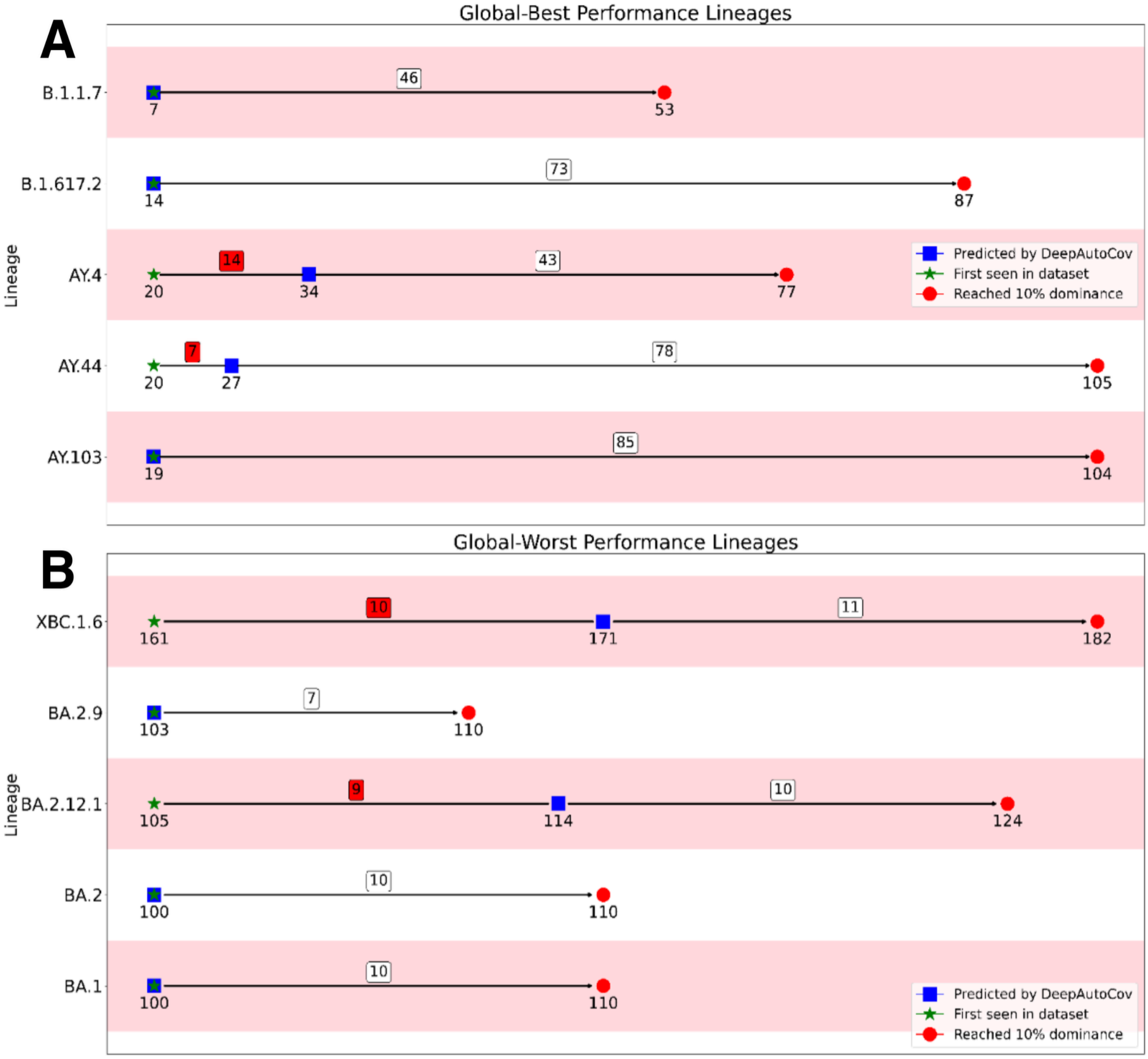

**Figure S3:** DeepAutoCoV best performances for FDL identification on the national data sets. The green star indicates the week when the FDL first appears in the data set. The blue square represents the week for first recognition. The red dot indicates the week when the FDL becomes dominant. **A)** USA, **B)** UK, **C)** Denmark, **D)** France.

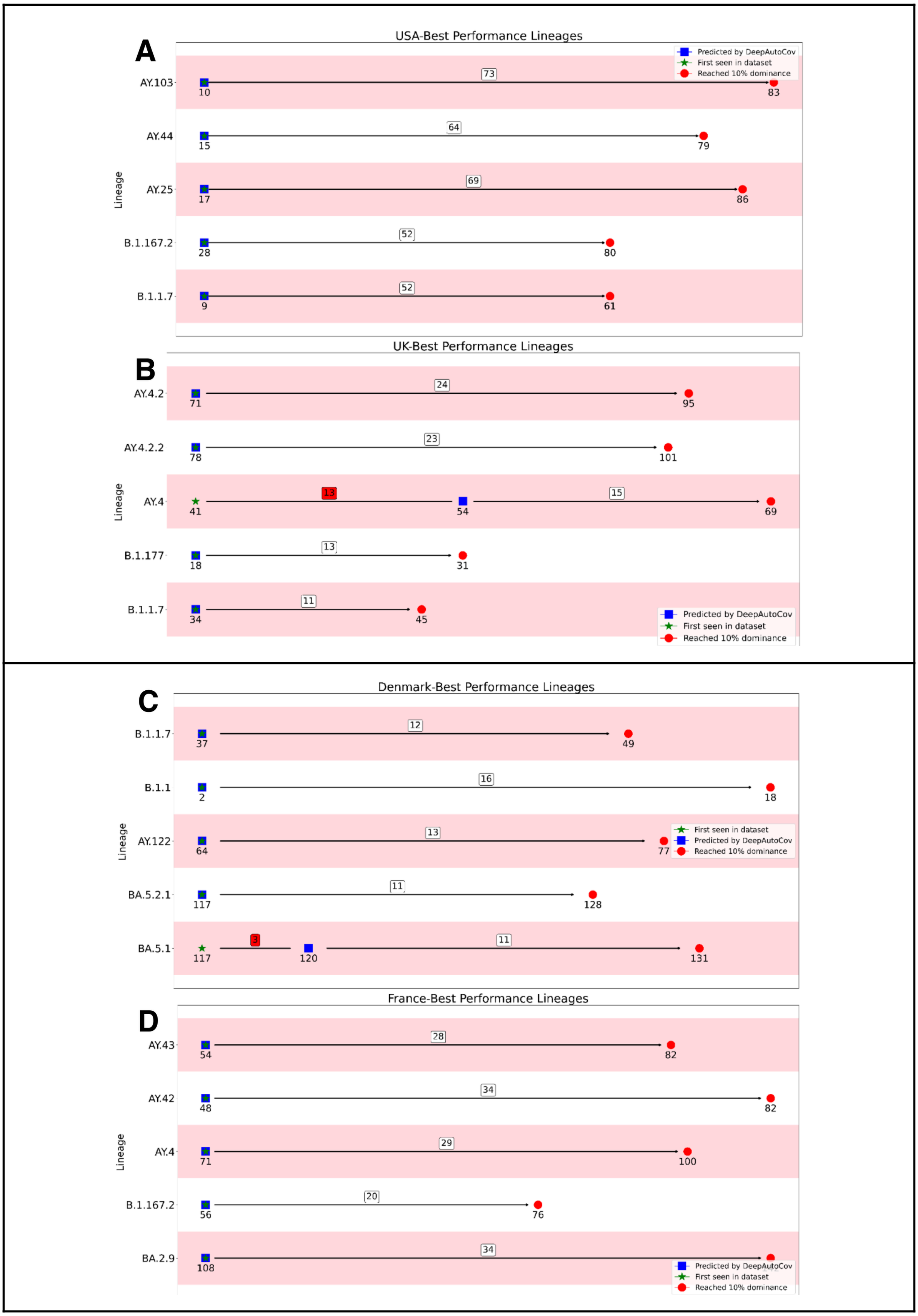

**Figure S4.** Top 10 FDLs correctly identified across different Data sets. We report the total number of FDLs (p-value)

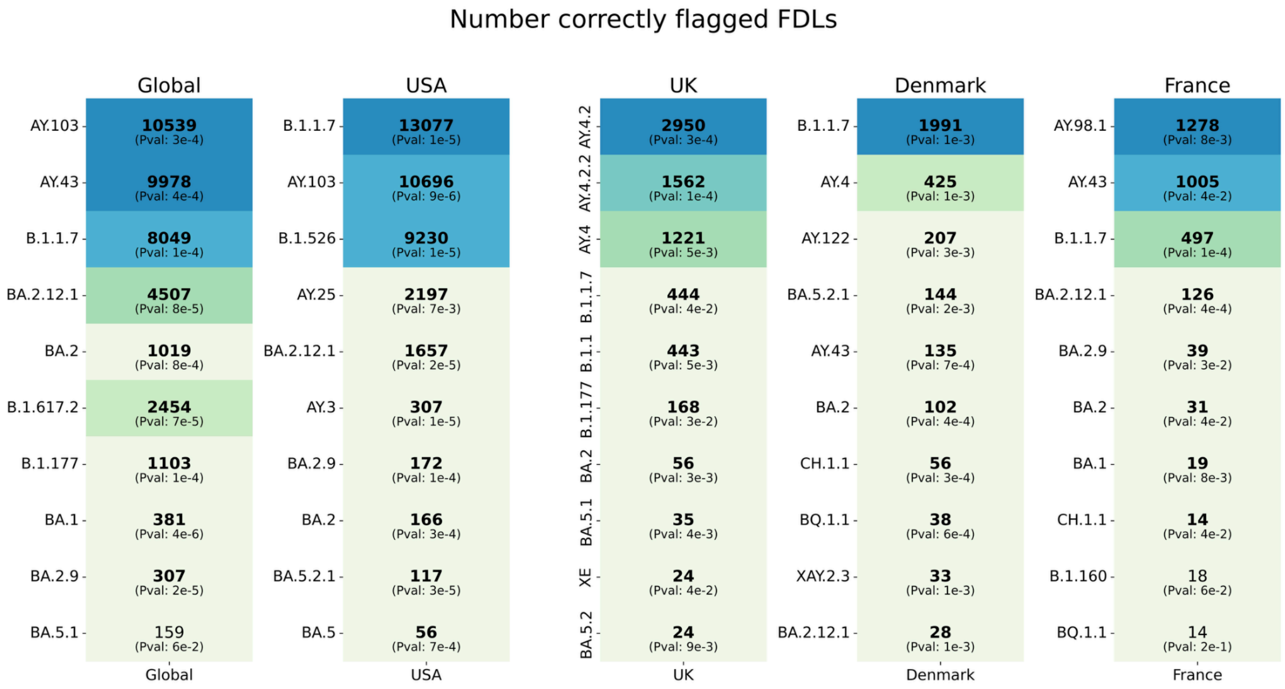

**Figure S5.** Top 10 Lineages identified as anomalies by DeepAutoCoV in 2024 ranking for MSE. Next to each bar appears the value of the PAM (DAYHOFF) Matrix score. The red line represents a MSE threshold used by DeepAutoCoV to to discriminate whether a sequence is anomalous or not

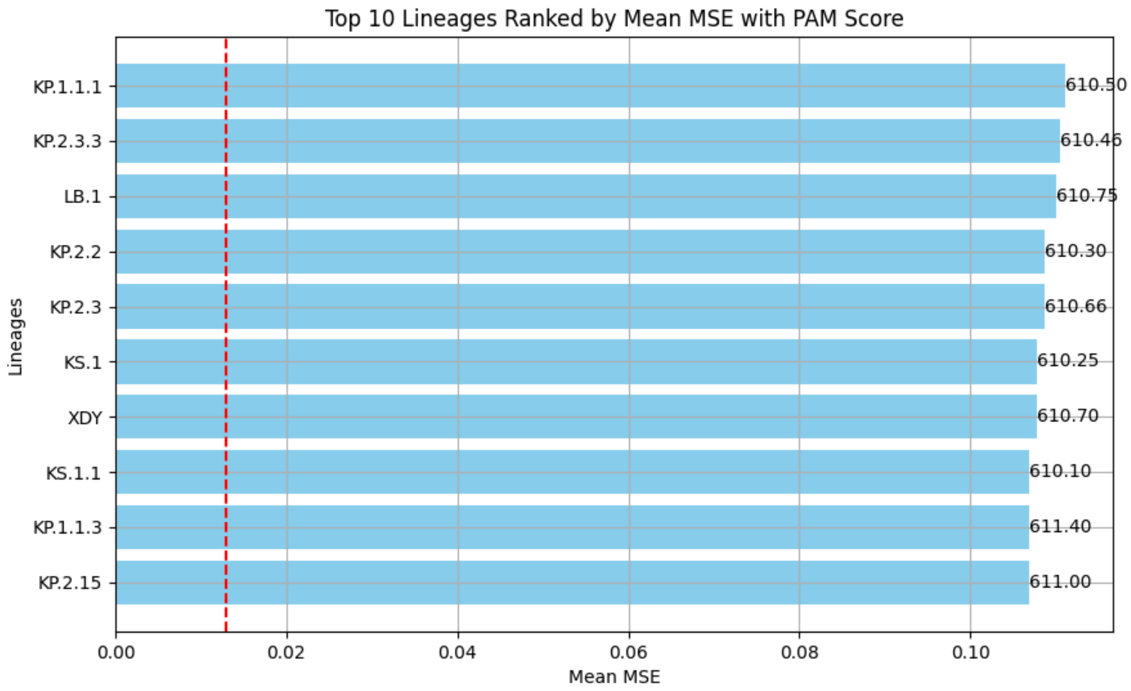
