## Supplementary Tables for "Forecasting dominance of SARS-CoV-2 lineages by anomaly detection using deep AutoEncoders"

**Table S1: Pre- and post-filtering total sequences for each of the data sets.**

| Data set | No. Spike sequences before filtering | No. Spike sequences after filtering |
|---|---|---|
| USA | 4,952,389 | 1,294,936 (27%) |
| UK | 3,118,879 | 962,496 (30%) |
| Denmark | 651,471 | 312,612 (48%) |
| France | 655,587 | 140,757 (23%) |
| Global | 16,187,950 | 4,506,983 (27%) |

SARS-CoV-2 Spike protein sequences sampled between December 24$^{th}$, 2019, and November 8$^{th}$, 2023, were downloaded from GISAID. **Filtering criteria:** human origin, full genome available, high coverage, without missing or incomplete submission date and without unknown amino acid (i.e. "X"), length within range (see Methods).

**Table S2: PPV comparison.**

| Method | Global | USA | UK | Denmark | France |
|---|---|---|---|---|---|
| DeepAutoCoV | 0.30 (0.06, 0.42) | 0.48 (0.10, 0. 72) | 0.57 (0.24, 0.92) | 0.49 (0.02, 0.90) | 0.47 (0.17, 0.70) |
| DB | 0.05 (0, 0.09) | 0.03 (0, 0.04) | 0* | 0.04 (0, 0.07) | 0.02 (0, 0.08) |
| LR | 0* | 0* | 0* | 0* | 0* |
| S2V NB, S2VT NB, S2S RESNET | 0* | 0* | 0* | 0* | 0* |

Median PPV (IQR) calculated over the entire time interval (2019-2023). DeepAutoCoV was compared to other unsupervised (DB = Distance-based linear), and supervised (LR = Logistic regression, S2V = Spike2Vec, S2S = Spike2Signal, NB = Naïve Bayes) methods.

**Table S3: FDLs identified in the Global data set.**

| Lineage | First appearance[1] | First flagging[2] | 10% threshold[3] |
|---|---|---|---|
| B.1 | 2 (1st week, Jan 2020) | 2 (1st week, Jan 2020, 9%) | 11 (2nd week, Mar 2020) |
| B.1.1 | 7 (2nd week, Feb 2020) | Unflagged | 12 (3rd week, Mar 2020) |
| B.1.2 | 4 (3rd week, Jan 2020) | Unflagged | 45 (1st week, Nov 2020) |
| B.1.177 | 12 (3rd week, Mar 2020) | 13 (4th week, Mar 2020, 0.5%) | 41 (1st week, Oct 2020) |
| B.1.1.7 | 7 (2nd week, Feb 2020) | 7 (2nd week, Feb 2020, 0.7%) | 53 (4th week, Dec 2020) |
| B.1.617.2 | 15(2nd week, Apr 2020) | 15 (2nd week, Apr 2020, 0.01%) | 87 (4th week, Aug 2021) |
| B.1.1.284 | 10 (1st week, Mar 2020) | 10 (1st week, Mar 2020, 1%) | 32 (1st week, Aug 2020) |
| AY.29 | 64 (2nd week, Mar 2021) | 64 (2nd week, Mar 2021, 0.01%) | 87 (2nd week, Sep 2021) |
| AY.4 | 34 (3rd week, Aug 2020) | 34 (3rd week, Aug 2020, 0.02%) | 77 (3rd week, Jun 2021) |
| AY.43 | 37 (2nd week, Sep 2020) | 37 (2nd week, Sep 2020, 0.02%) | 105 (4th week, Dec 2021) |
| AY.44 | 27 (2nd week, Jul 2020) | 27 (2nd week, Jul 2020, 0.03%) | 105 (4th week, Dec 2021) |
| AY.103 | 19 (2nd week, May 2020) | 19(2nd week May 2020, 0.02%) | 104 (3rd week, Dec 2021) |
| BA.1 | 100 (3rd week, Nov 2021) | 100 (3rd week Nov 2021, 0.01%) | 110 (3rd week, Jan 2022) |
| BA.2 | 100 (3rd week, Nov 2021) | 100 (3rd week Nov 2021, 0.03%) | 110 (3rd week, Jan 2022) |
| BA.2.9 | 103 (2nd week, Dec 2021) | 103 (2nd week Dec 2021, 0.03%) | 110 (3rd week, Jan 2022) |
| BA.2.12.1 | 105 (4th week, Nov 2021) | 114 (3rd week, Feb 2021, 0.05%) | 124 (2nd week, May 2022) |
| BA.5.1 | 121 (3rd week, Apr 2022) | 121 (3rd week, Apr 2022, 0.02%) | 137 (3rd week, Aug 2022) |
| BA.5 | 121 (3rd week, Apr 2022) | 121 (3rd week, Apr 2022, 0.08%) | 137 (3rd week, Aug 2022) |
| BA.5.2 | 123 (1st week, May 2022) | 127 (1st week, May 2022, 0.05%) | 140 (2nd week, Sep 2022) |
| XBB.1.5 | 140 (2nd week, Sep 2022) | 140 (2nd week, Sep 2022, 0.1%) | 163 (4th week, Feb 2023) |
| BR.2.1 | 141 (3rd week, Sep 2022) | 141 (3rd week, Sep 2022, 0.1%) | 153 (4th week, Dec 2022) |
| CH.1.1 | 144 (2nd week, Oct 2022) | 144 (2nd week, Oct 2022, 0.3%) | 167 (4th week, Mar 2023) |
| FK.1.1 | 164 (1st week, Mar 2023) | 164 (1st week, Mar 2023, 0.1%) | 182 (4th week, Jun 2023) |
| XBC.1.6 | 173 (2nd week, Feb 2023) | 173 (3rd week, Mar 2023, 1%) | 182 (4th week, Jun 2023) |
| XBC.1.3 | 142 (1st week, Oct 2022) | 142 (1st week, Oct 2022, 0.3%) | 185 (3rd week, Jul 2023) |
| DV.7.1 | 181 (3rd week, Jun 2023) | 181 (3rd week, Jun 2023, 1%) | 192 (2nd week, Sep 2023) |

| | | | |
|---|---|---|---|
| HW.1.1 | 181 ($3^{rd}$ week, Jun 2023) | 181 ($3^{rd}$ week, Jun 2023, 1%) | 195 ($1^{st}$ week, Oct 2023) |
| EG.5.1 | 180 ($2^{nd}$ week, Jun 2023) | Unflagged | 197 ($3^{rd}$ week, Oct 2023) |

Highlighted variants: yellow, Alpha; purple, Delta (sub)lineages; green, Omicron (sub)lineages. 1. Week number for FDL first appearance. 2. Week number for FDL first detection (frequency). 3. Week number for FDL reaching the threshold and becoming dominant.

**Table S4: FDLs identified in the USA data set.**

| FDLs | First appearance[1] | First flagging[2] | 10% threshold[3] |
|---|---|---|---|
| B.1 | 8 (4th week, Feb 2020) | 11 (3rd week, Mar 2020, 12%) | 11 (3rd week, Mar 2020) |
| B.1.2 | 2 (2nd week, Jan 2020) | 12 (4th week, Mar 2020, 1.2%) | 29 (3ht week, Jul 2020) |
| B.1.429 | 4 (4th week, Jan 2020) | 4 (4th week, Jan 2020, 7%) | 54 (2nd week, Jan 2021) |
| B.1.1.7 | 9 (1st week, Mar 2020) | 9 (1st week, Mar 2020, 0.2%) | 61 (2nd week, Mar 2021) |
| B.1.526 | 16 (2nd week, Apr 2020) | 16 (2nd week, Apr 2020, 0.1%) | 64 (1st week, Apr 2021) |
| AY.3 | 35 (1st week, Sep 2020) | 35 (1st week, Sep 2020, 0.04%) | 86 (3rd week, Sep 2021) |
| AY.25 | 17 (3rd week, Apr 2020) | 17 (3rd week, Apr 2020, 0.04%) | 86 (3rd week, Sep 2021) |
| AY.103 | 11 (3rd week, Mar 2020) | 11 (3rd week, Mar 2020, 0.06%) | 83 (4th week, Aug 2021) |
| AY.44 | 15 (1st week, Apr 2020) | 15 (1st week, Apr 2020, 0.04%) | 79 (4th week, Jul 2021) |
| B.1.617.2 | 28 (2nd week, Jul 2020) | 28 (2nd week, Jul 2020, 0.2%) | 80 (1st week, Aug 2021) |
| BA.1 | 102 (2nd week, Dec 2021) | 102 (2nd week, Dec 2021, 0.05%) | 110 (2nd week, Feb 2022) |
| BA.2.9 | 105 (1st week, Jan 2022) | 108 (4th week, Jan 2022, 0.2%) | 117(1st week, Apr 2022) |
| BA.2 | 105 (1st week, Jan 2022) | 105 (2nd week, Jan 2022, 0.04%) | 112 (4th week, Feb 2022) |
| BA.2.3 | 105 (1st week, Jan 2022) | 109 (1st week, Feb 2022, 1%) | 117(1st week, Apr 2022) |
| BA.2.12.1 | 102 (2nd week, Dec 2021) | 102 (2nd week, Dec 2021, 0.03%) | 120 (4th week, Apr 2022) |
| BA.5.2.1 | 127 (1st week, May 2022) | 127 (1st week, May 2022, 0.1%) | 136 (4th week, Aug 2022) |
| BA.5 | 128 (2nd week, Jun 2022) | 128 (2nd week, Jun 2022, 0.3%) | 136 (4th week, Aug 2022) |
| BA.5.1 | 127 (3rd week, Jun 2022) | 127 (3rd week, Jun 2022, 0.6%) | 136 (4th week, Aug 2022) |
| BA.5.2 | 127 (3rd week, Jun 2022) | 129 (1st week, Jul 2022, 0.1%) | 136 (4th week, Aug 2022) |
| BQ.1.1 | 142 (1st week, Oct 2022) | 142 (1st week, Oct 2022, 1%) | 157 (1st week, Jan 2023) |
| XBB.1.5 | 138 (1st week, Sep 2022) | 138 (1st week, Sep 2022, 0.8%) | 162 (2nd week, Feb 2023) |
| CH.1.1 | 144 (3rd week, Oct 2022) | 144 (3rd week, Oct 2022, 0.8%) | 169 (3rd week, Apr 2023) |
| FK.1.2.2 | 169 (3rd week, Apr 2023) | 169 (3rd week, Apr 2023, 4%) | 170 (4th week, Apr 2023) |
| FK.1.5 | 169 (3rd week, Apr 2023) | 176 (2nd week, Jun 2023, 11%) | 176 (2nd week, Jun 2023) |
| DV.1.1 | 164 (3rd week, Apr 2023) | 164 (3rd week, Apr 2023, 1%) | 176 (2nd week, Jun 2023) |
| FK.1.1 | 167 (2nd week, Apr 2023) | 167 (2nd week, Apr 2023, 3%) | 181(3rd week, Jul 2023) |
| XBB.1.16 | 180 (2nd week, Jul 2023) | 184 (2nd week, Aug 2023, 3%) | 188 (2nd week, Sep 2023) |
| DV.7.1 | 187 (1st week, Sep 2023) | 187 (1st week, Sep 2023, 2%) | 193 (2nd week, Oct 2023) |

Highlighted variants: yellow, Alpha; purple, Delta (sub)lineages; green, Omicron (sub)lineages. 1. Week number for FDL first appearance. 2. Week number for FDL first detection (frequency). 3. Week number for FDL reaching the threshold and becoming dominant.

**Table S5: FDLs identified in the UK data set.**

| FDLs | First appearance[1] | First flagging[2] | 10% threshold[3] |
|---|---|---|---|
| B.1.177 | 18 (3rd week, May 2020) | 18 (3rd week, May 2020, 0.3%) | 31 (3rd week, Aug 2020) |
| B.1.1.7 | 34 (4th week, Aug 2020) | 34 (4th week, Aug 2020, 0.05%) | 45 (1st week, Nov 2020) |
| B.1.1 | 3 (3rd week, Feb 2020) | 5 (1st week, Mar 2020, 8%) | 6 (2nd week, Mar 2020) |
| B.1 | 4 (4th week, Feb 2020) | 4 (4th week, Feb 2020, 7%) | 6 (4th week, Mar 2020) |
| AD.2 | 13 (2nd week, Apr 2020) | 13 (2nd week, Apr 2020,0.1%) | 31 (3rd week, Aug 2020) |
| AY.4 | 54 (3rd week, Jan 2020) | 54 (3rd week, Jan 2020, 0.01%) | 69 (3rd week, Mar 2021) |
| AY.4.2 | 71 (2nd week, Apr 2021) | 71 (2nd week, Apr 2021, 0.1%) | 95 (1st week, Oct 2021) |
| AY.4.2.2 | 78 (2nd week, Jun 2021) | 79 (2nd week, Jun 2021, 0.01%) | 101 (1st week, Nov 2021) |
| BA.2 | 103 (3rd week, Nov 2021) | 103 (2nd week, Nov 2021, 0.4%) | 108 (2nd week, Feb 2022) |
| XE | 104 (4th week, Jan 2022) | 104 (4th week, Jan 2022, 5%) | 108 (4th week, Feb 2022) |
| BA.5.2.1 | 128 (2nd week, Jun 2022) | 128 (2nd week, Jun 2022, 9%) | 132 (2nd week, Aug 2022) |
| BA.1 | 101 (1st week, Nov 2021) | 101 (1st week, Nov 2021, 0.02%) | 108 (2nd week, Feb 2022) |
| BA.2.12.1 | 119 (2nd week, Apr 2022) | 119 (2nd week, Apr 2022, 0.8%) | 123 (2nd week, May 2022) |
| BA.2.75 | 126 (1st week, Jun 2022) | 126 (1st week, Jun 2022, 4%) | 131 (1st week, Aug 2022) |
| BA.2.76 | 129 (4th week, Jun 2022) | 129 (4th week, Jun 2022, 8%) | 131 (1st week, Aug 2022) |
| BN.1 | 140 (2nd week, Oct 2022) | 140 (2nd week, Oct 2022, 8%) | 142 (4th week, Oct 2022) |
| CH.1.1 | 145 (1st week, Dec 2022) | 145 (1st week, Dec 2022, 9%) | 147 (3rd week, Dec 2023) |
| DV.6 | 155 (1st week, Feb 2023) | 155 (1st week, Jan 2023,2%) | 161 (4th week, Mar 2023) |
| DV.7 | 159 (2nd week, Mar 2023) | 159 (2nd week, Mar 2023, 2%) | 168 (2nd week, May 2023) |
| XBB.1.5 | 172 (2nd week, Jun 2023) | 172 (2nd week, Jun 2023, 9%) | 174 (4th week, Jun 2023) |
| XBB.1.16 | 172 (2nd week, Jun 2023) | 172 (2nd week, Jun 2023, 9%) | 174 (4th week, Jun 2023) |
| EG.5.1 | 177 (3rd week, Jul 2023) | 177 (3rd week, Jul 2023, 8%) | 183 (1st week, Sep 2023) |
| DV.7.1 | 187 (1st week, Oct 2023) | 187 (1st week, Oct 2023, 9%) | 190 (4th week, Oct 2023) |
| EG.5.1.3 | 182 (4th week, Aug 2023) | 182 (4th week, Aug 2023, 8%) | 190 (4th week, Oct 2023) |

Highlighted variants: yellow, Alpha; purple, Delta (sub)lineages; green, Omicron (sub)lineages. 1. Week number for FDL first appearance. 2. Week number for FDL first detection (frequency). 3. Week number for FDL reaching the threshold and becoming dominant.

**Table S6: FDLs identified in the DENMARK data set.**

| FDLs | First appearance[1] | First flagging[2] | 10% threshold[3] |
|---|---|---|---|
| B.1.177 | 26 (2nd week, Aug 2020) | 26 (2nd week, Aug 2020, 4%) | 29 (1st week, Sep 2020) |
| B.1.1 | 2(2nd week, Feb 2020) | 3 (3rd week, Feb 2020, 4%) | 18 (4th week, Jul 2020) |
| B.1.160 | 22 (2nd week, Aug 2020) | 26 (1st week, Sep 2020, 0.2%) | 29 (1st week, Sep 2020) |
| B.1.1.7 | 37 (1st week, Nov 2020) | 37 (1st week, Nov 2020, 0.4%) | 49 (1st week, Feb 2021) |
| AY.43 | 67 (4th week, Apr 2021) | 71(4rd week, May 2021, 3%) | 75 (4th week, Jun 2021) |
| AY.4 | 67 (4th week, Apr 2021) | 67 (4th week, Apr 2021, 0.2%) | 73 (2nd week, Jun 2021) |
| AY.122 | 67 (4th week, Apr 2021) | 67 (4th week, Apr 2021, 0.4%) | 75 (4th week, Jun 2021) |
| AY.7.1 | 68 (2nd week, May 2021) | 72 (1st week, Jun 2021,0.3%) | 75 (4th week, Jun 2021) |
| AY.4.6 | 75 (4th week, Jun 2021) | 75 (4th week, Jun 2021, 0.3%) | 81 (2nd week, Aug 2021) |
| BA.2 | 94 (3rd week, Nov 2021) | 95 (4st week, Nov 2021, 0.1%) | 98 (3rd week, Dec 2021) |
| BA.2.9 | 94 (3rd week, Nov 2021) | 95 (4st week, Nov 2021, 0.07%) | 98 (3rd week, Dec 2021) |
| BA.2.12.1 | 111 (4th week, Mar 2022) | 113 (2nd week, Apr 2022, 0.8%) | 120 (1st week, Jun 2022) |
| BA.5 | 117 (2nd week, Mar 2022) | 121 (1st week, Jun 2022, 0.2%) | 128 (4th week, Jul 2022) |
| BA.5.1 | 121 (1st week, Jun 2022 | 121 (1st week, Jun 2022, 0.1%) | 131 (3rd week, Aug 2022) |
| BA.5.2.1 | 113 (2nd week, Feb 2022) | 113 (2nd week, Feb 2022, 0.03%) | 128 (4th week, Jul 2022) |
| BF.7 | 121 (2nd week, Jun 2022) | 121 (2nd week, Jun 2022, 0.2%) | 130 (2nd week, Aug 2022) |
| BQ.1.1 | 135 (3rd week, Sep 2022) | 135 (3rd week, Sep 2022, 0.05%) | 148 (1st week, Jan 2023) |
| CH.1.1 | 143 (1st week, Oct 2022) | 143 (3st week, Oct 2022, 0.9%) | 155 (4th week, Feb 2023) |
| XBB.1.5 | 148 (1st week, Jan 2023) | 148 (1st week, Jan 2023, 0.6%) | 160 (1st week, Apr 2023) |
| FK.1.2.1 | 155 (4th week, Feb 2023) | 161(2n week, Apr 2023, 5%) | 161(2n week, Apr 2023) |
| DV.6 | 150 (3rd week, Jan 2023) | 150 (3rd week, Jan 2023, 1%) | 163 (4th week, Apr 2023) |
| EG.5 | 181 (2nd week, Sep 2023) | 181 (2nd week, Sep 2023, 30%) | 181 (2nd week, Sep 2023) |
| DV.7.1 | 181 (2nd week, Sep 2023) | 181(2nd week, Sep 2023, 4%) | 187 (1st week, Nov 2023) |

Highlighted variants: yellow, Alpha; purple, Delta (sub)lineages; green, Omicron (sub)lineages. 1. Week number for FDL first appearance. 2. Week number for FDL first detection (frequency). 3. Week number for FDL reaching the threshold and becoming dominant.

| FDLs | First appearance [1] | Firs flagging [2] | 10% threshold [3] |
|---|---|---|---|
| B.1.160 | 2 (2nd week, Jan 2020) | 2 (2nd week, Jan 2020, 3%) | 28 (2nd week, Jul 2020) |
| B.1 | 8 (3rd week, Feb 2020) | 4 (4th week, Feb 2020, 7%) | 13 (4th week, Mar 2020) |
| B.1.367 | 30 (1st week, Aug 2020) | 30 (1st week, Aug 2020, 6%) | 34 (4th week, Aug 2020) |
| B.1.177 | 18 (1st week, May 2020) | 18 (1st week, May 2020, 0.5%) | 45 (1st week, Nov 2020) |
| B.1.1.7 | 45 (2nd week, Nov 2020) | 45 (2nd week, Nov 2020, 0.3%) | 55 (3rd week, Jan 2021) |
| B.1.617.2 | 56 (4st week, Jan 2021) | 56 (4st week, Jan 2021, 0.5%) | 76 (3rd week, May 2021) |
| AY.98.1 | 72 (3rd week, Apr 2021) | 72 (3rd week, Jun 2021, 0.2%) | 84 (4th week, Jul 2021) |
| AY.42 | 48 (4th week, Nov 2022) | 48 (4th week, Nov 2022, 0.04%) | 82 (2nd week, Jul 2021) |
| AY.43 | 53 (1st week, Jan 2021) | 53 (1st week, Jan 2021, 0.2%) | 82 (2nd week, Jul 2021) |
| AY.4 | 70 (1st week, Apr 2021) | 70 (1st week, Apr 2021, 0.6%) | 100 (2nd week, Dec 2021) |
| AY.5 | 75 (2nd week, May 2021) | 80 (3rd week, Jun 2021, 0.08%) | 100 (2nd week, Dec 2021) |
| AY.122 | 70 (1st week, Apr 2021) | 78 (1st week, Jun 2021, 1%) | 94 (4th week, Oct 2021) |
| AY.125 | 78 (3rd week, Jun 2020) | 79(3rd week, Jun 2021, 1%) | 94 (4th week, Oct 2021) |
| BA.2 | 105 (3rd week, Feb 2022) | 105 (3rd week, Feb 2022, 0.2%) | 112 (2nd week, Apr 2022) |
| BA.2.12.1 | 123 (1st week, Jun 2022) | 124(2nd week, Jun 2022, 0.8%) | 130 (1st week, Sept 2022) |
| BA.2.9 | 108 (2nd week, Mar 2022) | 108 (2nd week, Mar 2022, 1%) | 142 (3rd week, Nov 2022) |
| BA.1 | 103 (1st week, Feb 2022) | 104 (2nd week, Feb 2022, 0.6%) | 110 (4th week, Mar 2022) |
| BA.2.75 | 135 (2nd week, Oct 2022) | 135 (2nd week, Oct 2022, 0.4%) | 144 (1st week, Dec 2022) |
| BA.5 | 123 (1st week, Jun 2022) | 123 (1st week, Jun 2022, 1%) | 138 (4th week, Oct 2022) |
| BE.6 | 142 (3rd week, Nov 2022) | 147 (3rd week, Dec 2022, 9%) | 148 (1st week, Jan 2023) |
| BF.7 | 135 (2nd week, Oct 2022) | 135 (2nd week, Oct 2022, 1.4%) | 148 (1st week, Jan 2023) |
| BQ.1.1 | 145 (2nd week, Dec 2023) | 145 (2nd week, Dec 2023, 0.05%) | 152 (1st week, Feb 2023) |
| BN.1 | 151 (4th week, Dec 2023) | 151 (4th week, Dec 2023, 4%) | 161 (3rd week, Apr 2023) |
| CH.1.1 | 151 (1st week, Feb 2022) | 151 (1st week, Feb 2022, 2%) | 159 (1st week, Apr 2023) |
| XBB.1.5 | 166 (4th week, May 2023) | 166 (4th week, May 2023, 9%) | 168(2nd week, Jun 2023) |
| XBB.1.16 | 176 (2nd week, Aug 2023) | 176 (2nd week, Aug 2023, 7%) | 188 (1st week, Nov 2023) |

**Table S7: FDLs identified in the FRANCE data set.**

Highlighted variants: yellow, Alpha; purple, Delta (sub)lineages; green, Omicron (sub)lineages. 1. Week number for FDL first appearance. 2. Week number for FDL first detection (frequency). 3. Week number for FDL reaching the threshold and becoming dominant.

**Table S8: (Sub)lineages detected in each data set during the 1st week (according to sampling date).**

| Region | 1st week sequencing | Variants in 1st week |
|---|---|---|
| USA | 04/01/2020 - 11/01/2020 | B, B.1.2, A |
| UK | 29/01/2020 - 6/02/2020 | A.2 |
| Denmark | 28/02/2020 - 6/03/2020 | B.1 |
| France | 01/01/2020 - 07/01/2020 | B.1.160 |
| Global | 24/12/2019 - 31/12/2019 | B |
